## Supplemental Material for "A cryptic pocket in Ebola VP35 allosterically controls RNA binding"

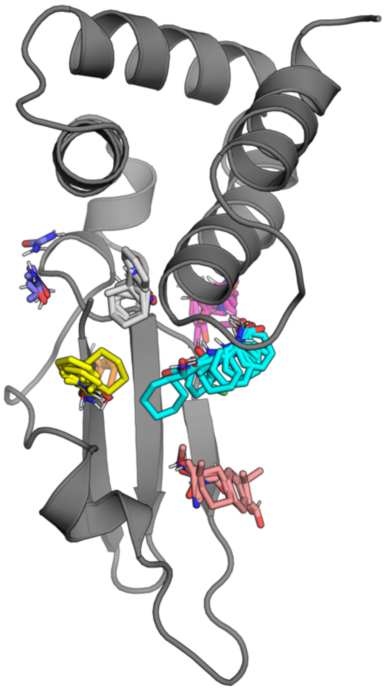

**Supplementary Figure 1.** FTMap results for the main cryptic pocket highlighting an example protein structure (gray) and hotspots where a variety of small organic probes (multicolored sticks) form energetically favorable interactions. The probe molecules are intended to capture different drug-like interactions (such as hydrogen bonding and Van der Waals contacts) and include acetamide, acetonitrile, acetone, acetaldehyde, methylamine, benzaldehyde, benzene, isobutanol, cyclohexane, N,N-dimethylformamide, dimethyl ether, ethanol, ethane, phenol, isopropanol, or urea.[1-4]

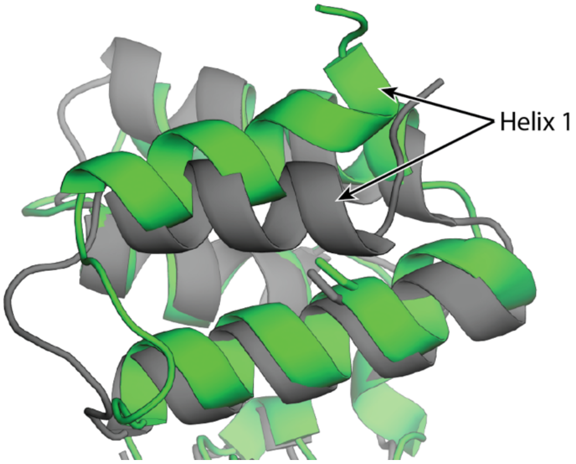


**Supplementary Figure 2.** Motion of helix 1 (green vs gray structures) sometimes exposes C247 (sticks) to solvent. However, the resulting pocket is small and FTMAP does not identify any hotspots in this region that are likely to bind drug-like molecules. Therefore, we focus our attention on the cryptic pocket created by the displacement of helix 5.

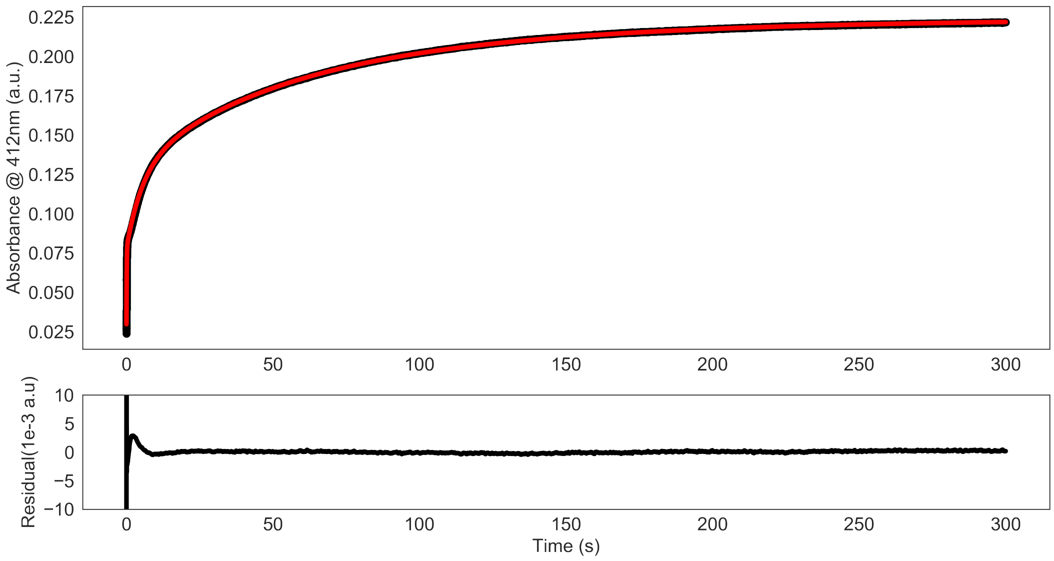


**Supplementary Figure 3.** A representative time trace from a thiol labeling experiment (black) performed at 100 μM DTNB and a quadruple exponential fit (red). The data are background subtracted (e.g. the average absorbance from three runs with DTNB but no protein were subtracted) to account for spontaneous hydrolysis of DTNB.

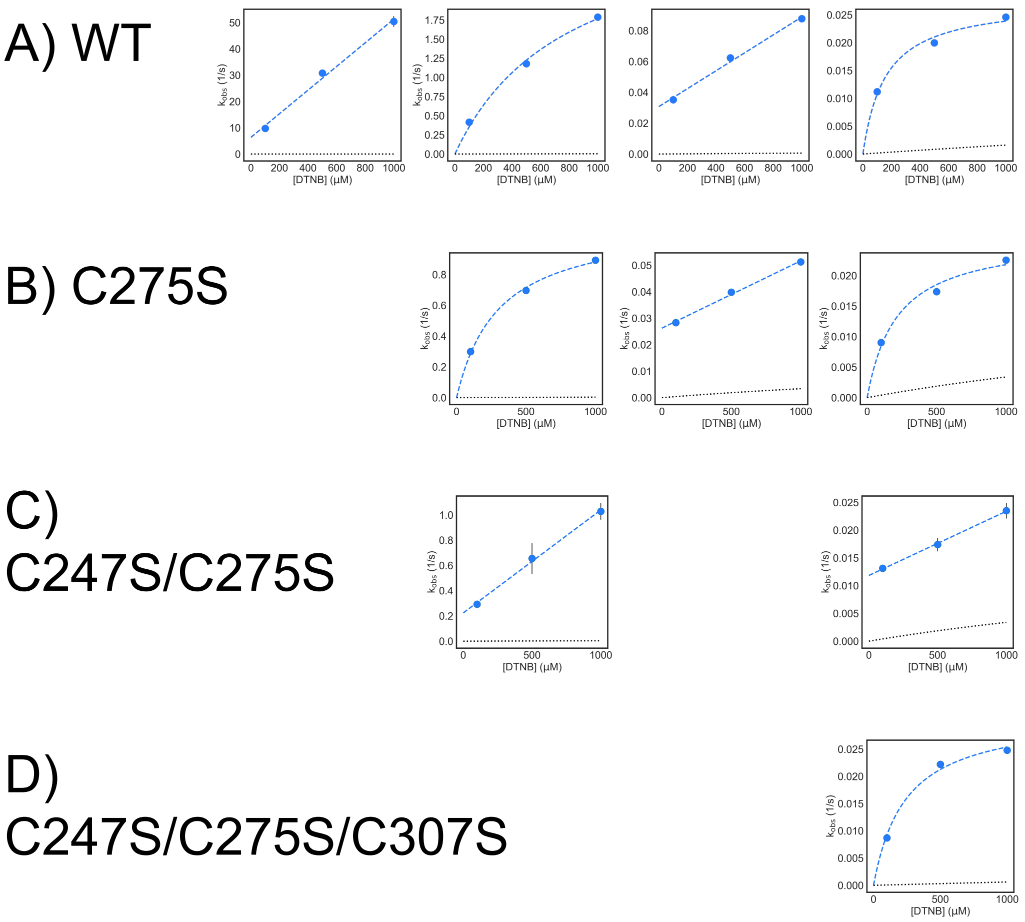


**Supplementary Figure 4.** Representative Kobs vs [DTNB] plots from thiol labeling experiments performed at 100. 500 and 1000 µM DTNB (blue circles) and fit of the Linderstrøm-Lang Model (dashed lines). A) wild-type observed rates. B) C275S observed rates C) C247S/C275S observed rates. D) C247S/C275S/C307S observed rates .

| Variant | C275 rate ($s^{-1}$) | C307 rate ($s^{-1}$) | C247 rate ($s^{-1}$) | C326 rate ($s^{-1}$) |
|---|---|---|---|---|
| Wild-type | 1.4 ± 0.02 | 0.201 ± 0.002 | 0.024 ± 0.0004 | 0.011 ± 0.000030 |
| C275S | - | 0.30 ± 0.002 | 0.028 ± 0.0003 | 0.0090 ± 0.00003 |
| C247S/C275S | - | 0.29 ± 0.01 | - | 0.013 ± 0.0004 |
| C247S/C275S/C307S | - | - | - | 0.0087 ± 0.0003 |
| C247S/C275S/C326S | - | 0.20 ± 0.007 | - | - |

**Supplementary Table 1.** Observed labeling rates at 100 μM DTNB for a set of variants with different cysteines mutated to serines to uncover which rate in the wild-type fit corresponds to which cysteine residue. Error is standard deviation from three replicates. Dash represents rates not measured due to the absence of that cysteine residue.

| Variant | K | $k_{unfold}$ ($s^{-1}$) |
|---|---|---|
| **Wild-type** | 6.57 ± 4.0 x $10^{-5}$ | 0.0175 |
| **C247S/C275S** | 4.01 ± 0.8 x $10^{-4}$ | 0.0083 |
| **F239A** | 4.67 ± 1.0x$10^{-4}$ | 0.0179 |
| **A291P** | 3.83 ± 4.0 x $10^{-5}$ | 0.00274 |

**Supplementary Table 2.** Characterization of the folding/unfolding of VP35's IID used to test whether the observed thiol labeling is due to fluctuations within the native state or global unfolding of the protein. K is the equilibrium constant between the folded and unfolded state determined from denaturation data, $k_{unfold}$ is the unfolding rate of the respective variants measured by intrinsic tryptophan fluorescence.

| Residue | $k_{int}$ ($\mu M^{-1}$ $s^{-1}$) |
|---|---|
| **C247** | 0.0566 ± 0.0007 |
| **C275** | 0.00254 ± 0.001 |
| **C307** | 0.0290 ± 0.002 |
| **C326** | 0.395 ± 0.02 |

**Supplementary Table 3.** Intrinsic labeling rates ($k_{int}$) for each cysteine residue. Intrinsic labeling rates were measured using either urea unfolded variants containing only the specified cysteine, or peptides containing the specified cysteine and its surrounding residues using an SX 20 stopped-flow uv-vis apparatus.

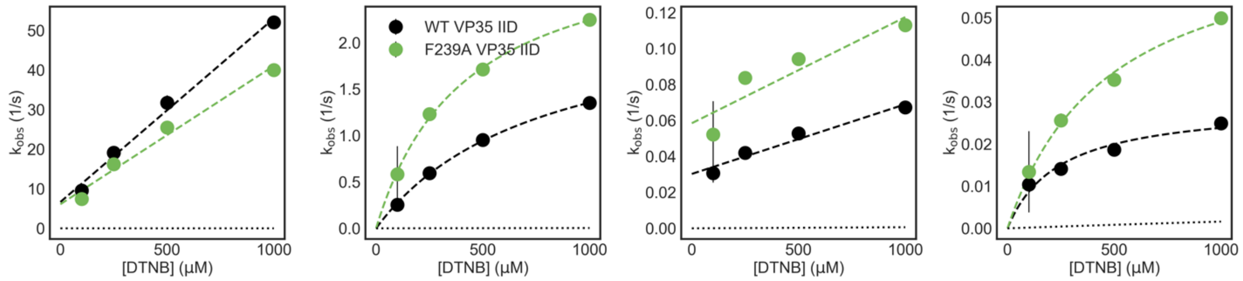


**Supplementary Figure 5.** Representative Kobs vs [DTNB] plots from thiol labeling experiments performed at 100, 250, 500 and 1000 µM DTNB for wild-type eVP35 (blue circles) and F239A (green circles) and fit of the Linderstrøm-Lang Model (dashed blue and green lines).

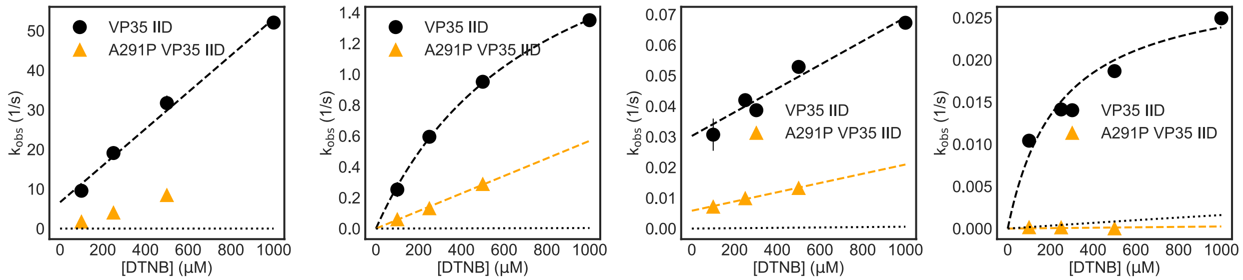


**Supplementary Figure 6.** Representative Kobs vs [DTNB] plots from thiol labeling experiments performed at 100, 250, 500 and 1000 μM DTNB for wild-type (black circles) and 100, 250, and 500 μM DTNB for A291P VP35 IID (orange circles) and fit of the Linderstrøm-Lang Model (dashed black and orange lines).

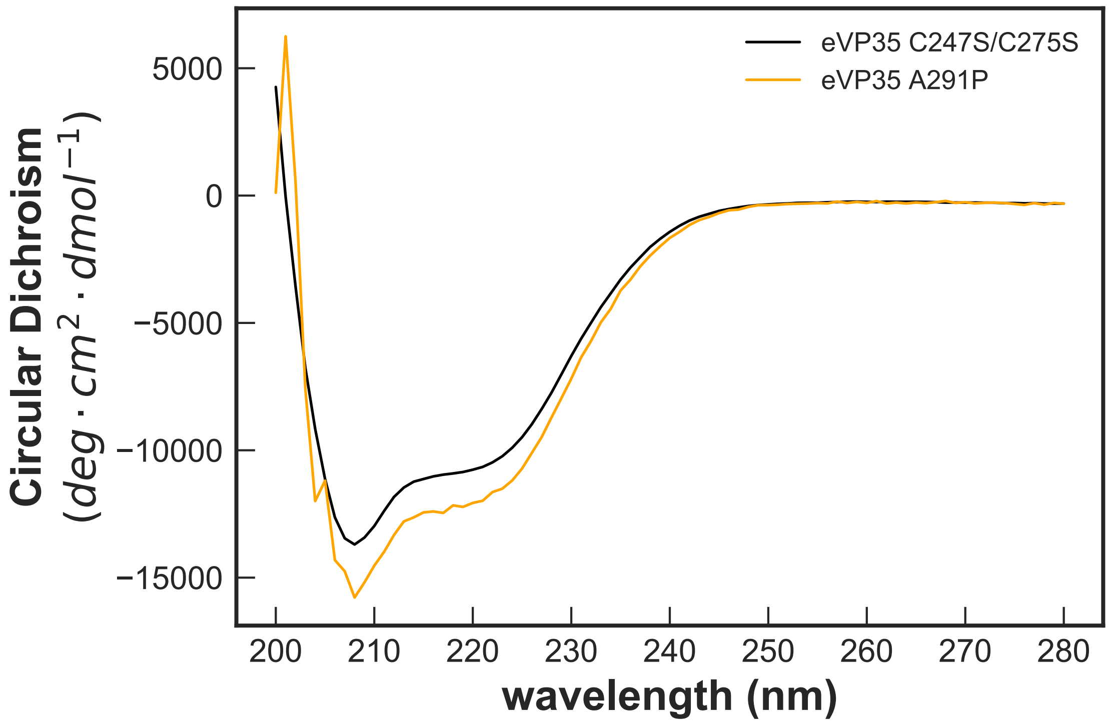


**Supplementary Figure 7.** Circular dichroism (CD) spectra of C247S/C275S (black) and A291P VP35 IID (orange) protein demonstrate that A291P substitution does not unfold the protein. The opaque and semi-transparent lines represent the mean and standard deviation, respectively, from three replicates. CD spectra were collected in 50 mM Sodium Phosphate pH 7 at 50 µg/mL protein (C247S/C275S) or 20 mM Tris pH 8, 150 mM NaCL at 35 µg/mL (A291P)

| Length | Sense Strand | Antisense Strand |
|---|---|---|
| 25mer | 56-FAM-rArArArCrUrGrArArArGrGrGrArGrArArGrUrGrArArArGrUrG | rCrArCrUrUrUrCrArCrUrUrCrUrCrCrCrUrUrUrCrArGrUrUrU |
| 25mer with 2nt 3' overhang | 56-FAM-rArArArCrUrGrArArArGrGrGrArGrArArGrUrGrArArArGrUrGrCrU | rCrArCrUrUrUrCrArCrUrUrCrUrCrCrCrUrUrUrCrArGrUrUrUrCrU |

**Supplementary Table 4.** RNA sequences used in fluorescence polarization binding assays. The sense and antisense strands were annealed in a 1:1 molar ratio.[5]

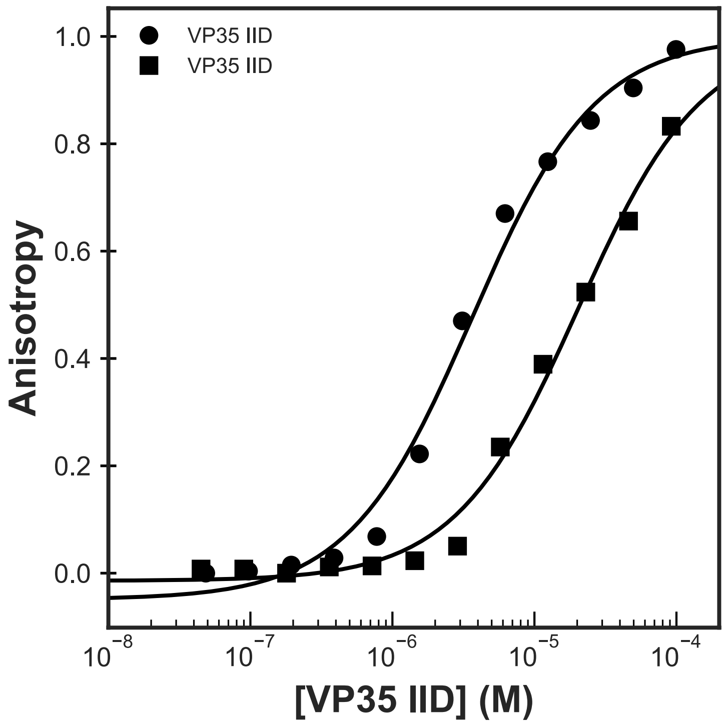


**Supplementary Figure 8.** Binding of C247S/C275S VP35 IID to a fluorescently labeled 25-bp double-stranded RNA with (squares) and without a two nucleotide overhang on the 3' end (circles). The anisotropy was calculated from measured fluorescence polarization and fit to a single-site binding model (black lines). The means and standard deviations from three replicates are shown but error bars are generally smaller than the symbols. Anisotropy was normalized to the max anisotropy for each data set.

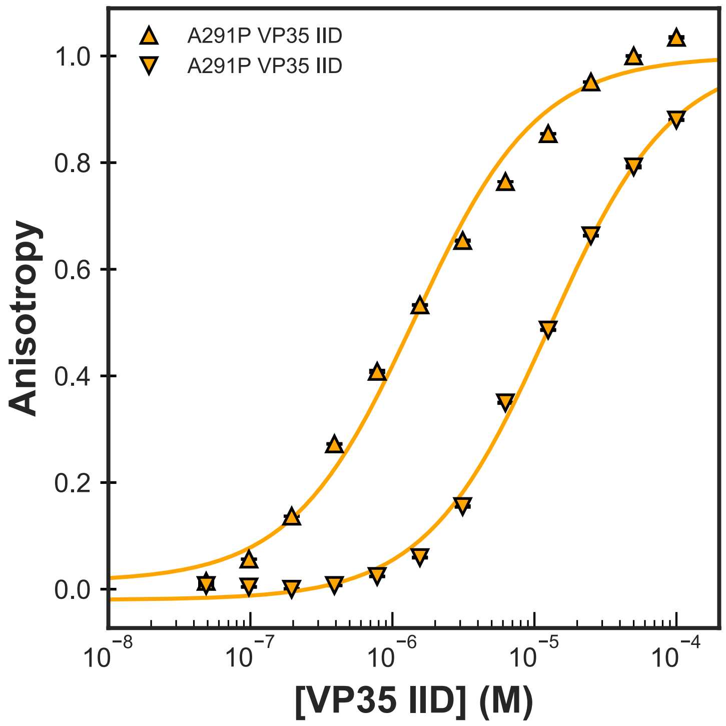


**Supplementary Figure 9.** Binding of A291P VP35 IID to a fluorescently labeled 25-bp double-stranded RNA with (squares) and without a two nucleotide overhang on the 3' end (circles). The anisotropy was calculated from measured fluorescence polarization and fit to a single-site binding model (black lines). The means and standard deviations from three replicates are shown but error bars are generally smaller than the symbols. Anisotropy was normalized to the max anisotropy for each data set.

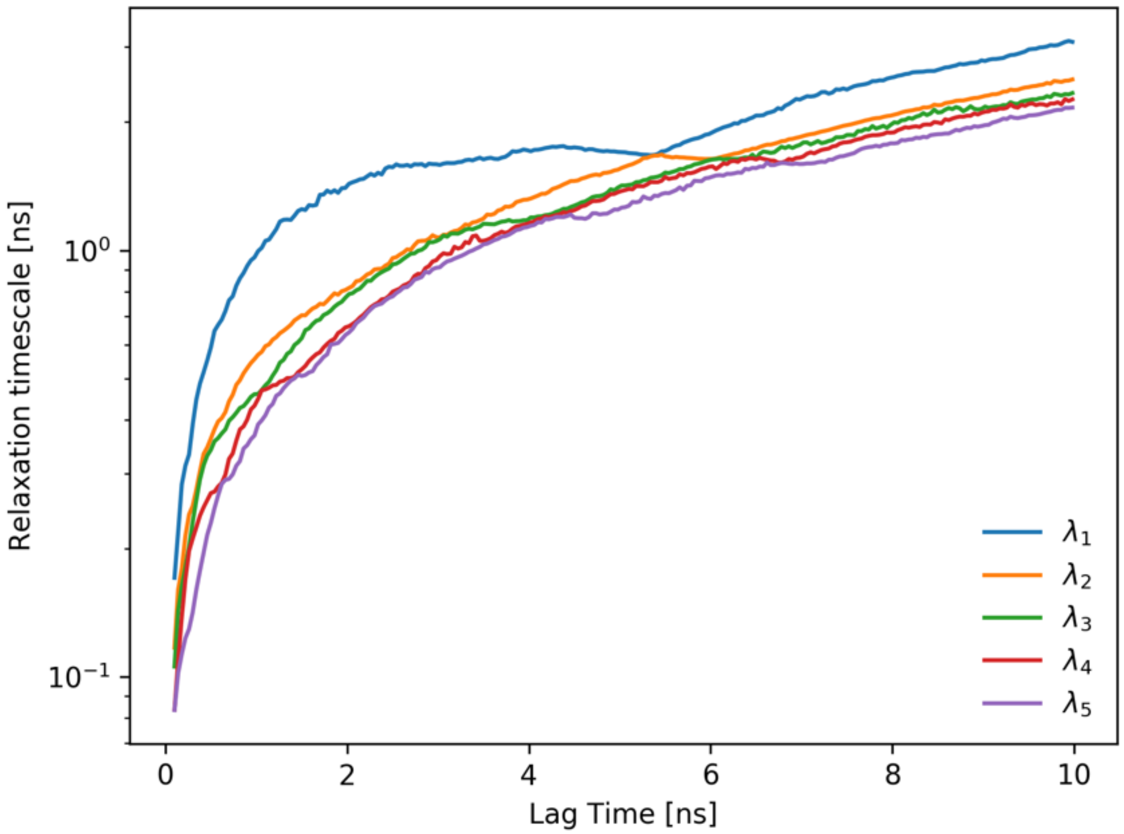


**Supplementary Figure 10.** Implied timescales test for the VP35 IID MSM suggests the kinetics are stable from 3-6ns. Analysis in the main text uses a Markov time of 6 ns. Key results were consistent for lag times from 3-6 ns.

**Supplemental Methods:**

We performed all data analysis and curve fitting using Python 3, Scipy and Numpy, in Jupyter Notebook. These notebooks are available upon request.

*Intrinsic Tryptophan Fluorescence Denaturation Experiments:*

To determine the denaturation midpoint and free energy of folding, eVP35 iid was equilibrated at 0.035 mg/mL in various concentrations of urea (MidSci, St. Louis, MO) from 0 to 8 M in 20 mM Tris pH 8, 150 mM NaCl. At 25ºC using a Jasco FP-8300 Spectrofluorometer the sample was excited with 280 nm light and we collected an emission spectrum from 300-400 nm. The max fluorescence emission at 0 M urea occurs at 322 nm. We recorded the fluorescence value at 322 nm for each concentration of urea in triplicate. Then, we used a six parameter fit for a two-state model of protein unfolding to determine the $C_M$ and folding free energy.[6] The folding free energy (ΔG) was used to calculate the fraction folded and fraction unfolded at 0 M urea. This was repeated for WT eVP35 iid, eVP35 iid C247S/C275S, eVP35 iid F239A, and eVP35 iid A291P.

$$Unfolded \overset{K}{\leftrightarrow} Native$$

$$Obs\ Fluor = \frac{Fluor_{U,H2O} + \beta_U[Urea] + \left(Fluor_{N,H20} + \beta_N[Urea]\right)e^{-\frac{(\Delta G_{H2O}+m[Urea])}{RT}}}{1 + e^{-\frac{(\Delta G_{H2O}+m[Urea])}{RT}}}$$

Fluor is the baseline fluorescence values for the native and unfolded states absent any denaturant.

β values are the slopes of the native and unfolded baseline signals.

To determine the unfolding rates, WT eVP35 iid, C247S/C275S, F239A, or A291P were manually injected into a solution of 20 mM Tris pH 8, 150 mM NaCl to a final concentration of 0.03 mg/mL at varying concentrations of urea equal to or above the $C_M$ measured in the equilibrium experiments. Then we collected fluorescence measurements exciting at 280 nm and detecting at 322 nm until the resulting curve reached steady state ~500s or more. The resulting curve was fit to an exponential model to extract the observed unfolding rate at that concentration of urea. Then we plotted the log of the observed rates as a function of urea concentration and linearly extrapolated to the unfolding rate in 0 M urea.

*Circular Dichroism Spectra:*

We evaluated the effect of amino acid substitutions on protein folding and overall secondary structure formation via circular dichroism spectrophotometry. Protein samples were buffer exchanged into the buffer at which they were measured and diluted to 50 or 35 µg/mL (see specific figure legends for variant specific information). We collected spectra at 25˚C in an Applied Photophysics Chirascan equipped with a Quantum Northwest Inc. TC125 Peltier-controlled cuvette holder, reading every integer wavelength between 200 and 280 nm, averaging data for each point for either one or five seconds. Data reported are in either millidegrees, or converted into molar ellipticity using the following equations[7].

$$\theta = (mDeg * MRW)/(10 * l * C)$$
$$MRW = M/(N - 1)$$

Here, θ is molar ellipticity, mDeg is millidegrees CD, MRW is the mean residual weight, $l$ is the path length in centimeters, C is the protein concentration in molar, M is the protein molecular weight in g/mol and N is the number of amino acids in the protein.

*RNA Fluorescence Anisotropy Experiments:*

Monitoring the binding of protein to 25-bp dsRNA gives affinities that are consistent with past work. Previous work used a dot-blot assay to measure binding and reported an apparent dissociation constant ($K_d$) for blunt-ended dsRNA of 3.4±0.07 μM.[5] Similarly, our FP assay gives an apparent $K_d$ of 3.64 ± 0.34 μM for VP35 IID binding blunt-ended dsRNA (Fig. 5B). Furthermore it was previously reported that the addition of 2-nucleotide overhang to the 3' end of the RNA reduces VP35 dsRNA-binding affinity by 10-fold.[8] Even when the presence of an overhang inhibits blunt end binding the VP35 IID is still able to bind to the dsRNA backbone though this interaction is weaker than with the blunt ends. Similarly to published data, our FP assay gives an apparent $K_d$ of 20.4±1.1 μM corresponding to at least a five fold reduction in apparent binding to the overhang dsRNA relative to the blunt ended dsRNA. However, an upper baseline could not be captured due to limitations in the protein's solubility, so this apparent $K_d$ is a lower bound. These data are also fit well assuming an apparent $K_d$ of 30.1±7.2 μM that was reported previously.[8] Monitoring the binding of VP35 to 25-bp dsRNA with two blunt ends or 3' overhangs demonstrates that our FP assay is sensitive to both dsRNA-binding modes and gives affinities that are consistent with past work (Fig. 5B).

We buffer exchanged and concentrated to ~300 μM VP35 iid then serially diluted by half across the rows of Costar 96 well half area black flat bottom polystyrene plates with a non-binding surface, in triplicate, in 10 mM Hepes pH 7, 150 mM NaCl, 1 mM $MgCl_2$. Then we added dsRNA to each well at a final concentration of 100 nM 25 bp FAM-dsRNA. For every experiment we included the following control wells: buffer only, dsRNA only, dsRNA with VP35 at the expected $K_D$, and dsRNA with VP35 and a previously described dsRNA binding inhibitor[9]. The plates were protected from light and allowed to equilibrate for at least one hour before reading as described in the main-text methods. Data were routinely checked for fluorescence anomalies in the control and experimental wells.


1 Kozakov, D. *et al.* Structural conservation of druggable hot spots in protein–protein interfaces. *Proceedings of the National Academy of Sciences* **108**, 13528-13533, doi:10.1073/pnas.1101835108 (2011).

2 Kozakov, D. *et al.* The FTMap family of web servers for determining and characterizing ligand-binding hot spots of proteins. *Nature Protocols* **10**, 733-755, doi:10.1038/nprot.2015.043 (2015).

3 Brenke, R., Kozakov, D., Chuang, G. Y., Beglov, D. & 2009. Fragment-based identification of druggable 'hot spots' of proteins using Fourier domain correlation techniques. *Bioinformatics* **25**,621-627 10.1093/bioinformatics/btp036 (2009).
4 Bohnuud, T., Beglov, D., Ngan, C. H., acids, B. Z. N. & 2012. Computational mapping reveals dramatic effect of Hoogsteen breathing on duplex DNA reactivity with formaldehyde. *Nucleic Acids Research* **40**, 7644-7652*,* doi:10.1093/nar/gks519 (2012).
5 Edwards, M. R. *et al.* Differential Regulation of Interferon Responses by Ebola and Marburg Virus VP35 Proteins. *Cell Reports* **14**, 1632-1640, doi:10.1016/j.celrep.2016.01.049 (2016).
6 Street, T. O., Courtemanche, N. & Barrick, D. in *Methods in Cell Biology* Vol. 84 295-325 (Academic Press, 2008).
7 Greenfield, N. J. Using circular dichroism spectra to estimate protein secondary structure. *Nature Protocols* **1**, 2876-2890, doi:10.1038/nprot.2006.202 (2006).
8 Ramanan, P. *et al.* Structural basis for Marburg virus VP35–mediated immune evasion mechanisms. *Proceedings of the National Academy of Sciences of the United States of America* **109**, 20661-20666, doi:10.1073/pnas.1213559109 (2012).
9 Glanzer, J. G. *et al.* In silico and in vitro methods to identify ebola virus VP35-dsRNA inhibitors. *Bioorganic & Medicinal Chemistry* **24**, 5388-5392, doi:10.1016/j.bmc.2016.08.065 (2016).
